# Adaptational lags during periods of environmental change

**DOI:** 10.1101/742916

**Authors:** Tom J. M. Van Dooren

**Affiliations:** CNRS - UMR 7618 Institute of Ecology and Environmental Sciences (iEES) Paris, Sorbonne University, Case 237, 4 place Jussieu, 75005 Paris, France

**Keywords:** Adaptation, adaptive control, climate change, evolutionary ecology, maladaptation, invasion analysis, pest management

## Abstract

Effects of climate change can be handled by means of mitigation and adaptation. In the biological sciences, adaptations are solutions which evolved when organisms needed to match an ecological challenge. Based on Adaptive Dynamics theory, a definition is proposed of adapted states and adaptational lags applicable during periods with environmental change of any speed. Adaptation can thus be studied when it emerges from complex eco-evolutionary processes or when targets for adaptation are not defined a priori. The approach is exemplified with a model for delayed germination in an annual plant. Plasticity and maternal effects are often presumed to be adaptive and added to the model to investigate lags in these modes of trait determination. Adaptational lags can change sign and to understand their dynamics, effects of trait space boundaries and characteristics of years with large numbers of recruits had to be considered. Adaptational lags can be crucial elements of adaptive control strategies for managed ecosystems. To demonstrate their practical relevance, examples from pest management show that evolutionary adaptation has been used to infer targets of control. Adaptational lags then serve as measures of the distance to the control target and become integral elements of strategies for adaptive pest population management.

## Introduction

Climate change generates massive challenges to our societies. Other species will show a range of responses to environmental change (Newman et al. 2011), and these can be to our own advantage or not. Initially, human policy to deal with climate change mostly focused on mitigating our interactions with the environment (Schipper 2006), by reducing the speed and magnitude of change or by dampening the effects of environmental changes. However, awareness has increased that human adaptation is necessary, merits attention and that it interacts with mitigation (Kane & Shogren 2000).

In biology, adaptation is a term which predates the development of evolutionary biology (Amundson 1996). It is not equivalent to the presence of an evolutionary response. Walsh and Lynch (2018) for example, a reference work on selection and evolutionary responses, hardly use the term and equate “adaptive” to “beneficial”. Fisher (1930) stated that adaptation concerns assessing the conformity of an organismal phenotype to an adapted phenotype determined from an imagined comparison of several potential environments and phenotypes. In this view, adaptation is not merely a description of an actual evolutionary process, but an investigation of the conformity of phenotypes with predictions obtained from considering a wider range of environments and phenotypes. Adaptation is believed to be important for the survival of populations, but more recent modelling also found the opposite: evolutionary suicide can occur when populations adapt (Gyllenberg & Parvinen 2001). Above all, assuming adaptation helps formulating and testing hypotheses. When we assume perfect adaptation to arrive at sharp predictions, this leads to hypotheses which are generally falsifiable.

In the context of human climate change responses, “adaptation” has become reframed over the years from the eco-evolutionary meaning applicable to any organism into a policy response (Schipper 2006); from describing a dynamics of (autonomously) evolving systems into a concept more in line with human adaptive systems control towards a given target (Tyukin 2011). This trend complicates the integration of evolutionary adaptation by biological systems into adaptation policies. A revised or re-clarified notion of evolutionary adaptation to climate change could address this.

The consensus is that most organisms currently respond to climate change with phenotypic plasticity (Gienapp & Merilä 2018), i.e., developmental changes that are often believed to be adaptive in response to altered environmental conditions (Merilä & Hendry 2014). The alternative which is usually considered, evolution and adaptation by changes in genotype frequencies, requires the presence of heritable genetic variation for the involved traits.

Therefore, most studies looking for ongoing adaptation in natural systems focused on demonstrating that phenotypic responses were due to changes in gene frequencies (Gienapp & Merilä 2018). However, evolutionary responses don’t need to move systematically in the direction where we expect that adaptation would be fastest or even where phenotypic changes are adaptive (Leimar 2009, Teplitsky et al. 2014). On top of that, the target of adaptation can move over time. This has usually been modelled by making the target an explicit time-dependent parameter (Lynch et al 1991), i.e., as if the changes in the target of adaptation are highly predictable or known. For situations where the targets of adaptation need to emerge from a study of eco-evolutionary processes across environments and phenotypes, it is unclear how to predict them precisely or how to capture their time patterns during climate change with a simple function.

Below, I propose a strategy to determine adaptations during climate change. Essential to the proposal is that most of the concepts applied were already developed in so-called Adaptive Dynamics approximations (Metz et al. 1996, Geritz et al. 1998) also known as evolutionary invasion analysis (Otto & Day 2007). A classical model for delayed germination in annual plants with a seed bank (Cohen 1966, Ellner 1997) is used as an example of an analysis of evolutionary adaptation and adaptational lags during environmental change. I then attempt to characterize the combination of evolutionary adaptation and adaptive control and give examples of managed ecosystems where predictions of evolutionary adaptation are relevant for their control and where adaptational lags are measures of the distance from the control target.

A general approach to determine adaptations during environmental change Organisms adapt to a range or set of situations encountered, hence adaptation always accounts for environmental variability (Fisher 1930, Levins 1968). When adapted states are determined, it is often assumed that these variable environments don’t show any lasting trends over time and occur with fixed probabilities. The time series of the variables describing them are then stationary (Tuljapurkar 1986, Metz et al. 1992, Coulson 2020). On the other hand, projections of environmental variables in periods of climate change involve systematically changing temperature and rainfall averages (Collins et al. 2013). We describe these environmental changes as time-dependent changes in the parameters of probability distributions of environmental variables (Figure 1A). These parameters determine how the mean and dispersion of environmental variables change over time. We usually assume that they change gradually within the same probability distribution family. The actual environmental history observed will thus be a realization from a stochastic process with time-dependent parameters (Fig. 1B). Each probability distribution per time point, say year, is specified by a set of parameter values and this distribution of environmental states represents a collection of situations to which organisms can adapt. For each given set of parameter values of the environment, we can therefore determine what the adaptative phenotypes are (Fig. 1C). The result for a period of environmental change is then a time series of adapted stated determined using a time series of parameters of the distributions of environmental variables (Fig. 1C). Adapted states can be determined using Adaptive Dynamics approximations (Metz et al. 1996, see also Abrams 2005, Methods Supplement), which assume separated ecological and evolutionary timescales and simple genetics in order to focus the analysis on the relationship between ecological environments and phenotypes (Dieckmann 1997). Adaptive Dynamics approximations have been used to generate new hypotheses, to predict evolutionary outcomes driven by ecological interactions (Dieckmann 1997) and to develop management strategies, for example for the management of epidemies during climate change (Sabelis and Metz 2002). For evolving populations where genotypes are not necessarily at an adapted state all the time, we can determine how far each individual in the population is from a predicted adapted state at each given point in time (Fig. 1D). For example, for an individual with a scalar phenotypic trait *x* and an adapted phenotypic state at time *t* which is *x*_*t*_^*^, the adaptational lag is *x*_*t*_^*^ - *x*.

**Figure one.**
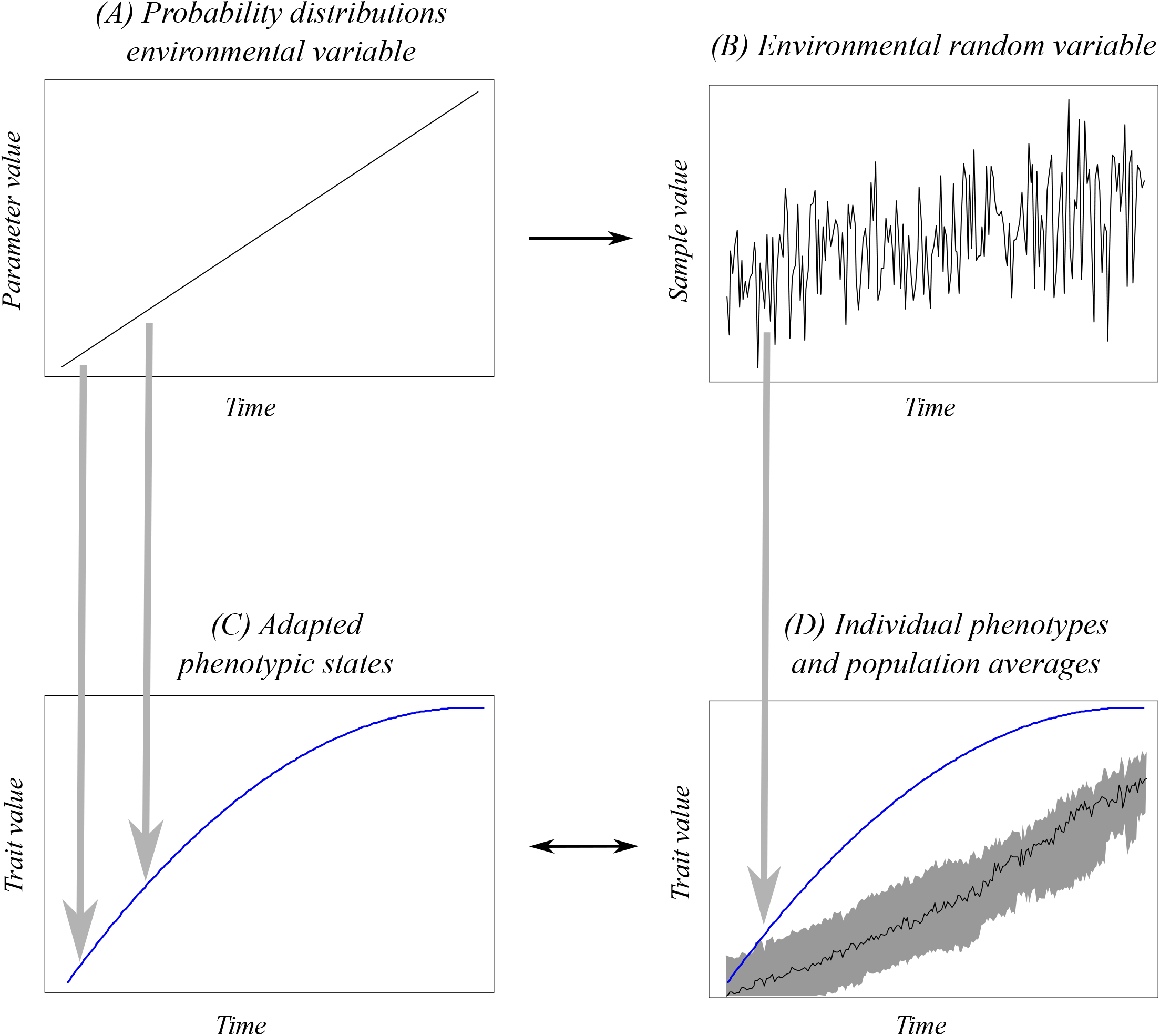
A general approach to determine adaptations and adaptational lags during periods of environmental change. (A) Environmental change is described by gradual changes in parameters of the probability distributions of environmental variables. (B) The actual environmental history is a single realization of this time-dependent stochastic process. (C) For each value or set of values of environmental parameters per time point, adapted states which evolution could arrive at can be determined (blue). (D) An actual population shows evolutionary responses depending on, inter alia, selection occurring at each point in time. At each point in time, there will be a range of phenotypes present (grey band), and an average phenotypic value across the population (black line). The difference between the phenotypic trait value or traits composing an adapted state and the phenotype of an individual is the adaptational lag. It therefore varies between individuals, and population statistics of this lag can be calculated.

### Adaptational lags in delayed germination

Several aspects of the determination of adaptational lags can be illustrated using a well-known model for the demography of an annual plant (Cohen 1966, Bulmer 1984). This model tracks densities (number per unit of area) of individual seeds of a single genotype in a seed bank across years, each time censused right after seed production. With *N*_*t*_ the number of seeds at the end of the season in year *t* and assuming large population sizes such that demographic stochasticity can be ignored, the number of seeds one year later is

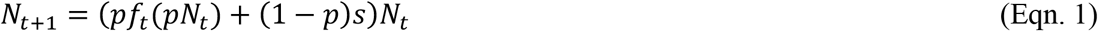

with *p* their probability of germination, *s* the survival probability in the seed bank across one year and *f*_t_ the per capita number of seeds contributed to the seed bank by a germinated seed in year *t*. The number of seeds recruited is stochastic and density-dependent. It depends on the total number germinated *pN*_t_. Across successive years the population grows in a multiplicative manner. The population density *N*_*t*+*τ*_ after an interval of *τ* years is:

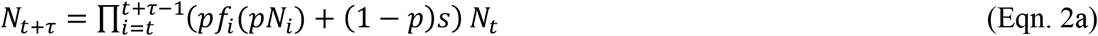

This is one of the simplest and best-known structured population models (during the growing season ungerminated seeds and plants are simultaneously present). It is the prime example used to demonstrate that adaptation is not always towards the fastest life history with the largest number of offspring (Stearns 2000).

This model is easily modified to allow for the occurrence of many genotypes in a population. We can then use simple forward simulations to investigate how selection gradually changes population composition among genotypes with different phenotypic traits. A common idea is that adapted states emerge when genotype frequencies equilibrate, for example by fixation of one genotype among several already present in a population. However, evolutionary invasion analysis (Adaptive Dynamics, Metz et al. 1996, Geritz et al. 1998, Methods Supplement) finds adapted states differently and also considers genotypes not yet present in a population. It values resistance to invasion and we therefore need to determine which genotypes can invade so-called resident populations (established populations, with stationary fluctuations or equilibria in their densities). More importantly, we must determine which resident population compositions (1) can evolve by means of a trait substitution sequence fuelled by mutations and (2) additionally have the property that once their state attained, they cannot be invaded by genotypes with similar germination phenotypes. Hence, they can persist in a longer-term evolutionary sense and are considered to be adapted to the ecological circumstances. In order to determine these states, invasion fitness (Metz et al. 1992) is used to assess the evolutionary stability (convergence stability and invasibility, see Methods Supplement) of resident population states. Here we assume that only a single resident phenotype affects the success of rare genotypes, and that the process characterising the resident state and the fluctuations in demographic parameters is stochastic, stationary and ergodic (Rand et al. 1994, Ferrière & Gatto 1995, Ripa & Dieckmann 2013) such that the precise starting point *t* becomes irrelevant. For a single resident genotype with the germination trait value *p* as above, and a rare mutant genotype with a different germination probability *p*’, invasion fitness equals

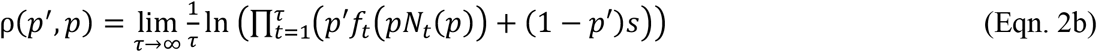

This expression assumes clonal inheritance and that the number of seeds recruited varies between years, even for an identical amount of germinated seeds. *N*_*t*_(*P*) is the number of seeds in the seed bank at time *t*, with the resident genotype having phenotype *p*. By extension, when survival in the seed bank is also allowed to be time-dependent, we get the invasion fitness

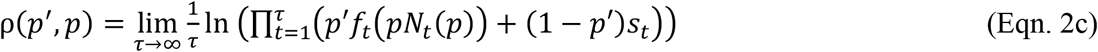

Invasion fitness is directly related to the establishment probability of a new strategy in a given stationary environment with a resident population already present (Haldane 1927, Ripa & Dieckmann 2013). Ellner (1997) contains an insightful presentation of results which exploit a partial derivative used to determine whether delayed germination, i.e., a germination probability below one, can invade. This derivative is 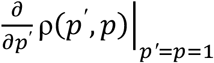 (Eqn. 3a). From its sign we know whether selection favours mutant germination probabilities smaller than the resident germination probability, which is equal to one. When it is positive, mutant germination probabilities below one are disfavoured and it is adaptive to always germinate. When it is negative, we can expect adapted states with a germination probability below one, i.e., the establishment of a seed bank where seeds remain one or several seasons. A crucial insight in evolutionary ecology was that such probabilities can only be adaptive when life history parameters fluctuate. In these cases, we need to determine the intermediate germination probability values that are so-called evolutionarily singular points (ESS, Geritz et al. 1998, represented here as *p*^*^). For resident populations with a single genotype, these are defined by the condition that the fitness derivative 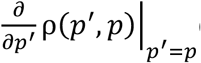 (Eqn. 3b) locally vanishes. There is no directional selection driving traits away from an ESS germination probability *p*^*^. Local second-order partial derivatives determine further requirements for adapted states: they need to be convergence stable, hence reachable, and non-invasible (see Methods). When adapted states consist of a single genotype, they are called CSS (Continuously Stable Strategy, Eshel 1983). There is no analytical expression for the adaptations *p*^*^. Their dependence on environmental parameters needs to be determined from numerical analysis.

## Continuation

If we assume that years can be good with probability *q* and otherwise bad, that the total recruitment to the seed bank equals *K* per unit of area in a good year and *k* in a bad year, then (Eqn. 3a) can be worked out. This is an exercise in Elmer (1997) which I don’t want to spoil, but the result is summarized in Figure two. For environments with fluctuations between good and bad years that occur according a Bernoulli distribution with probability *q*, it is adaptive to delay germination when recruitment parameters *K* and *k* of good and bad years differ sufficiently, when survival in the seed bank *s* is high and when probability *q* is intermediate (Fig. 2). Now arises the question what this tells us on adaptation when the environment changes in a non-stationary manner, for example when *s* and *q* would gradually change between years. A first line of attack could be to assume “continuation” (Kuznetsov 2004), namely that the population will always be able to attain the adapted state very rapidly and that gradual environmental changes provoke gradually changing population states that are each predicted well by the adapted state for the environment of that year (e.g., Ferrière & Legendre 2013). For the example model here, the state of the environment in a specific year is given by the four parameters *s, q, K* and *k*. Three examples of continuation trajectories are given in Fig. 2. The first two are for gradual changes in survival and the probability that years are good, with contrasting patterns. It can be first adaptive to have a seed bank when survival in the seed bank is still high, and when survival has decreased, to always germinate. The opposite route can also be adaptive: first always germinate, but when good years occur less often and survival gradually decreases, it can become adaptive to develop a seed bank. If recruitment parameters gradually change as well, differences between good and bad years could first become smaller and subsequently larger again. This scenario is elaborated below. However, Fig. 2 already shows that in such cases having a seed bank can first be adaptive, then it can become adaptive to always germinate, followed by a second period where delayed germination is adaptive.

**Figure two.**
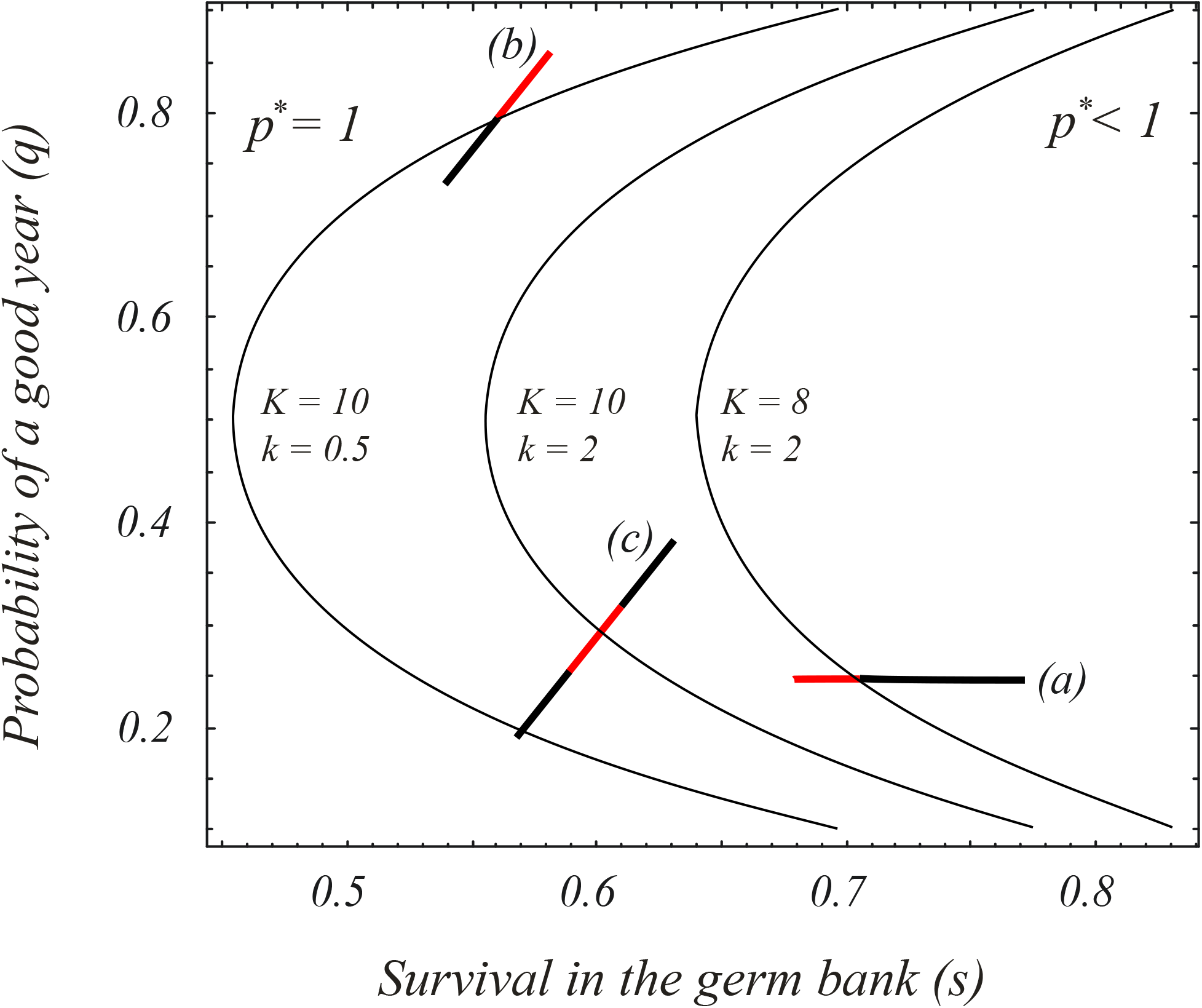
Continuation of adaptations during environmental change. We can determine which combinations of total recruitment in good (*K*) and bad years (*k*), survival in the seed bank (*s*) and the probability that a year is good (*q*) lead to (CSS) adaptations where the adaptive germination probability *p*^*\**^ is smaller than one. To the right of each curve labeled with the corresponding recruitment parameters (recruitment boundary), adapted germination probabilities for these combinations of *s* and *q* and the recruitment values specified on the boundary are smaller than one, hence a seed bank adaptive. When the difference between *K* and *k* increases, boundaries shift to the left and more combinations of *s* and *q* have a CSS *p*^*\**^ < 1. When environments change gradually, we can assume that parameter values (*s, q, k, K*) change gradually between years and that evolution tracks the adapted germination probability value specific to the environmental regime of each year. Three examples are given, with adapted states where *p*^*\**^ < 1 in black, with *p*^*\**^ = 1 in red: (a) If survival in the seed bank is decreasing, then it will become adaptive to always germinate. (b) Environmental change decreases *q* and *s* such that it is first adaptive to always germinate, later it becomes adaptive to develop a seed bank. (c) If recruitment parameters would gradually change as well, we can imagine that differences between good and bad years become smaller and then larger again, as in the scenario elaborated further in the main text.

## Adaptational lag

However, it is a strong assumption to make that populations would be well adapted all the time when the environment changes. For an individual with a genotypically determined germination probability *p* and an adapted state at time *t* which is *p*_*t*_^*^ the adaptational lag equals the difference *p*_*t*_^*^ - *p*. It is positive when the adapted state is a larger germination probability than the one for this individual, negative when it is smaller. The average adaptational lag in a population is the average of this quantity among the individuals present at a given census point. Even with high levels of standing additive genetic variation, non-zero average lags are predicted to occur when adapted states are boundary values and genetic variation remains present due to mutation pressure or segregation. In other cases, average adaptational lags are immediately generated when distributions of standing variation are asymmetric around an adapted state, when selection fluctuates over time or when organisms have complex life histories inherently generating lags in selection responses.

From now on, think of good and bad years as referring to initially warm and cool years (Figure 3), such that *q* is the probability of a warm year, *K* is the total recruitment in a warm year and *k* in a cold year. To explore potential time trajectories of adaptational lags, we investigate a scenario where the probability *q* of a relatively warm year gradually changes. At the same time, and mimicking an effect where all years become gradually warmer, *K* and *k* change in values over time (as in example C in Fig.2). When warm years become hot, *K* will decrease. At the same time, cold years gradually improve such that recruitment *k* increases. Survival probability in the seed bank gradually decreases, for example due to gradually increasing soil temperatures (Ooij 2012). This scenario has an intermittent period where an egg bank is not adaptive (Fig. 3). I determined the adapted state for each year in the scenario and simulated a population of selfing annual plants with a substantial mutation rate per individual (0.01 per individual, Methods, Fig. 3). The lag remains substantial even with this large continual input of new mutational variation. It is not the case that the population equilibrates at a constant lag distance from the CSS adaptive state as in many other models (Kopp and Matuszewski 2015). While the CSS moves towards a boundary trait value, remains there and subsequently moves away from it, the adaptational lag changes sign (Fig. 3). Mutational pressure on the germination probability near *p* = 1, makes it that the seed bank never completely disappears, even when that would be adaptive. This opposes the idea that increased amounts of heritable genetic variation will inherently lead to improved adaptation.

**Figure three.**
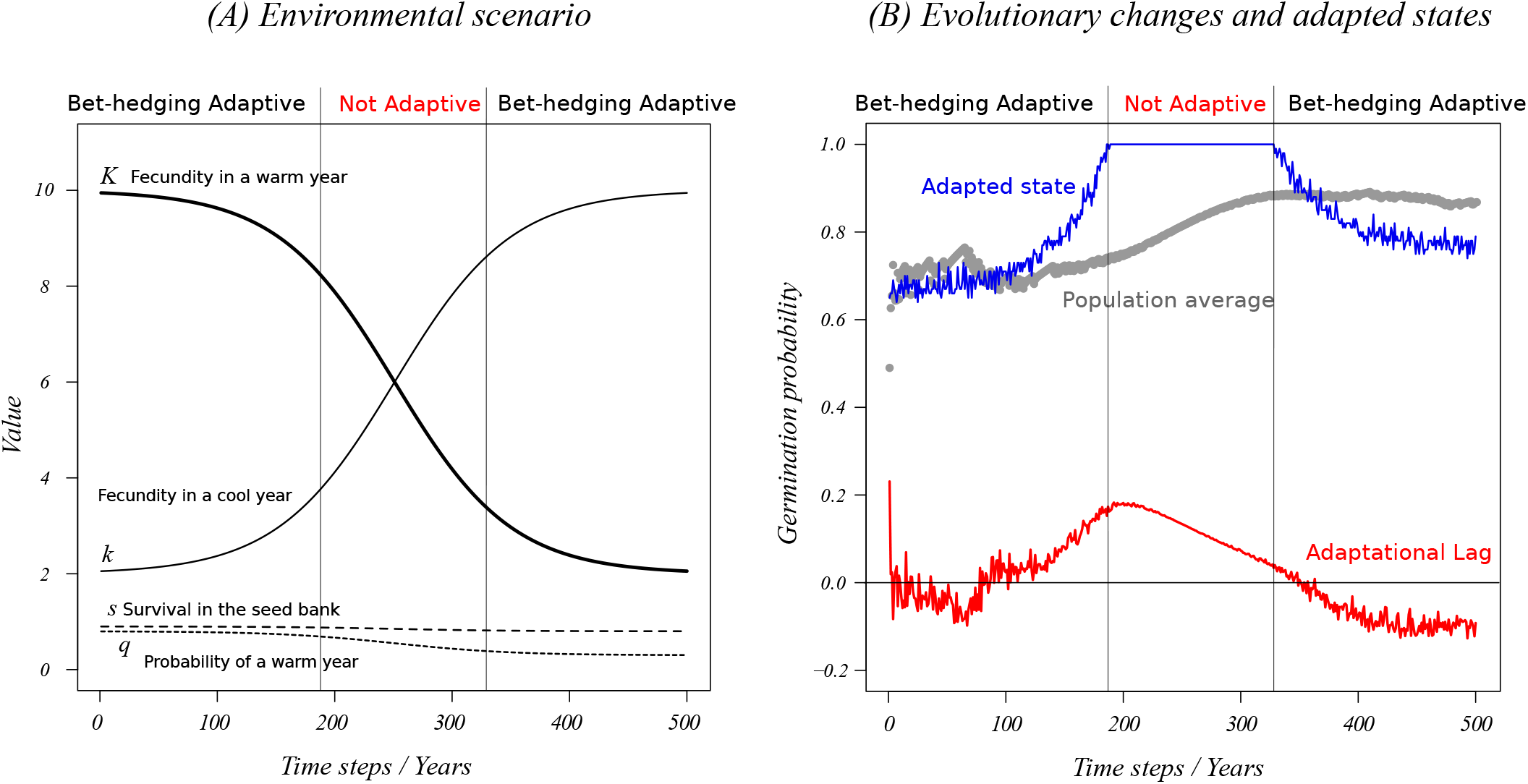
Adaptational lags for a single evolving trait. (A) A scenario of gradual change in the four parameters characterising the environment per year. As in the example (C) of Fig.2, the difference between recruitment parameters *K* (warm year) and *k* (cool year) first gradually decreases, then increases again. It can be determined that there is a time window from years 188 to 329 where it is adaptive to germinate with probability *p*^*\**^ = 1. (B) Calculation of the adaptational lag for the environmental change scenario in the left panel and a simulation of a population of selfing individuals with a high mutation rate in the germination probability. Full blue line: Adapted states determined using Adaptive Dynamics. Grey line: population trajectory, average germination probability of the seeds in the seed bank. Red line: difference between adapted states and population average.

### Phenotypic plasticity and parental effects

Additional relevant components of trait determination were added to this model, by allowing germination to depend on an environmental cue perceived by a plant embryo (phenotypic plasticity) or by the mother plant (a parental effect). Botero et al. (2015) already investigated environmental changes that demanded restructuring of phenotype determination, but did not work out any adaptational lags. For the annual plant example, the reaction norm specification becomes as follows (Eqn. 4):

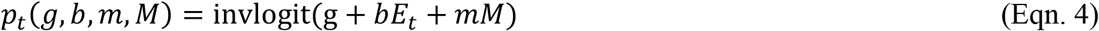

where invlogit denotes the inverse-logit function, invlogi*t*(*x*) *= e*^*x*^*/*(1 + *e*^*x*^).

Germination probability *P*_*t*_ now depends in a non-linear manner on a genotypic component of an individual *g*, on an environmental cue *E*_*t*_ and on a maternal cue *M* stored by that individual. The maternal and the environmental cues are weighed by specific factors, *b* and *m* respectively. These factors and the genotypic component *g* can evolve by mutations and selection. For the maternal cue, it is assumed that it is a marker left by a mother on an offspring individual, that it can have two states and that it is fixed and stored by the individual in the same state for the rest of its life. I assume that only part of the individuals are responsive to a cue for a good year or get the right cue from their parent. In the simulations presented, 70% of individuals respond to a warm year (warm year *E*_t_ = 1, cool year *E*_t_ = 0). In a warm year, 70% of individuals get the maternal cue *M* = 1, the others get *M* = 0. In a cool year, 70% of individuals get *M* = 0, the others *M* = 1. Because of the inverse-logit transformation, trait values *g, b, m* can range from minus to plus infinity, while the individual germination probabilities remain constrained between zero and one. Note that each individual has five state variables, of which three are evolving and adapting via mutations and selection. What is also important is that with maternal effects, the seed bank contains individuals in different states, namely those that carry maternal cue *M = 0* and the others carrying *M = 1*.

If we rerun the environmental scenario of Fig. 3 and determine the adapted CSS states per year for the three trait components, then adapted states for the plasticity weight are generally extreme (Figure 4A), while a pattern similar to Fig. 3 is observed for the genotypic value, with a shorter time interval where bet-hedging is not adaptive. The adaptive maternal effect weight is often opposite in sign to the plasticity weight. In fact, in part of the time interval where bet-hedging is not adaptive, all three traits jointly have their adapted states at maximum values. The eco-evolutionary dynamics of the simulated population shows a rather different pattern (Fig. 4B): the maximum average genotypic value *g* in the simulation is only reached when bet-hedging is already adaptive again. Note that the genotypic value *g* slightly increases further in the second time interval where bet-hedging is adaptive, even when its adaptational lag has already changed sign. The plasticity weight also evolves for part of the years in a direction opposite from where the adapted state is for these years. When adapted states are not determined by means of trait substitution sequences but approximated by simulated evolution in the environment characteristic for a year, the mutational pressures near trait boundaries and standing amounts of variation have effects on the outcome (Fig. 4D).

**Figure four:**
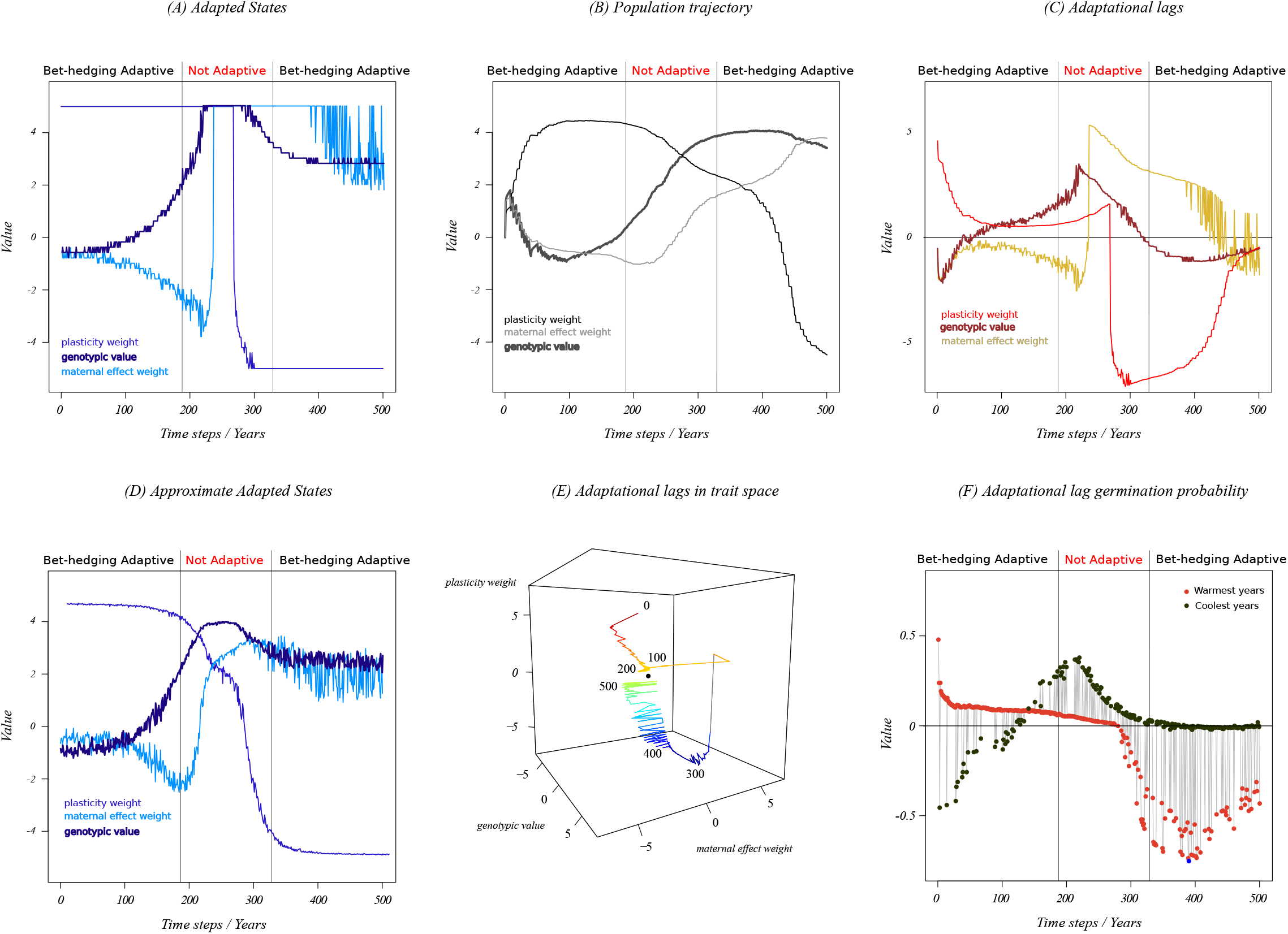
Adapted states and adaptational lags for a model version with multiple components of trait determination. The same environmental change scenario as in Fig. 2 is simulated. (A) Adapted states of the three trait components. The genotypic component is drawn in dark blue, the plasticity weight in blue and the maternal effect weight in light blue (B) Population trajectories of the averages of each trait component. (C) Average adaptational lag trajectories per trait component. (D) Approximate adapted states determined by evolution with a large supply of mutational variation in each stationary environment. (E) Average adaptational lag as a three-dimensional trajectory. (F) Average adaptational lag calculated for the germination probability as described by Eqn. 5. Points for warm and cool years are plotted in red and brown, respectively.

Approximate adapted states on trait space boundaries are much less observed than in Fig. 4A, and when a trait such as *g* has not evolved to the boundary, the parental effect for example seems to converge to a different value as well. This demonstrates that approximating adapted states by evolutionary simulations with a continual input of a lot of genetic variation is imprecise.

### Multivariate lags?

When individual phenotypes depend on several underlying component traits, we can calculate adaptational lags in different ways. It is straightforward to represent lags by vectors in a multivariate trait space, but there is no obvious standardization of each axis to make different trait components directly comparable. As Fisher (1930) focused on organismal individual phenotypes, we can focus on the germination phenotypes and substitute the evolving components of individuals in the actual population (*g, b, m*) by the values at the adapted state (*g*^*\**^, *b*^*\**^, *m*^*\**^), leaving the non-evolving state variables of each individual untouched (*E*_*t*_ and *M*). The adaptational lag of an individual then becomes

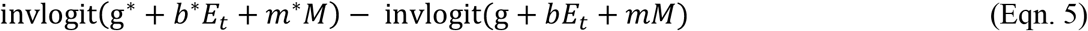

Note that a study of lags could provide evidence of genetic conflicts: selective pressures on heritable phenotype components evidenced by speeds of trait change, can change in dependence on the magnitude of the lag in others. Next to that, the average lags in the more elaborate model version with evolving genotypic values, plasticity and parental effects show non-trivial trajectories (Fig. 4E), with overshoots around an adapted state and average lag trajectories resembling those of a stable focus in other dynamical systems (Kuznetsov 2004).

Adaptational lag averages in the population can be calculated for warm and cool years separately, and this demonstrates (Fig. 4F) that in the first half of the simulation, lags become smallest in warm years, when these occur more often than cool years. In the second half, the lags become smallest in cool years. This suggests that evolution reduces adaptational lags in the most common environments fastest.

### Responsiveness to selection and establishment probabilities

The average adaptational lag can be useful as a measure of the lag in other population statistics. For example, many evolutionary models find the maximum population density at the adaptive state and the adaptational lag then represents the distance from that situation. In the model example it is less obvious what the practical relevance of the lag might be and this requires further exploration. Such a simple exploration was carried out for the model version with plasticity and maternal effects.

As a measure of turnover indicating how fast a population can respond to selection on juvenile and adult traits, I calculated per year which fraction of the seed bank at the end of the reproductive season was recruited in that same year. Figure four shows that it relates well to the average germination probability in a population (Fig. 5A), but that the dependence on the adaptational lag is complex and gives the lag little predictive value for this measure of responsiveness to selection (Fig. 5B). Peischl and Kirkpatrick (2012) found that establishment probabilities of populations in arbitrarily changing environments depend on the time pattern of fluctuations and that the per capita number of offspring of a new strategy in its first year has the largest weight on the outcome. When the log per capita number of offspring of germinated seeds is estimated, this relates poorly to the average germination probability in the population (Fig. 5C), while the adaptational lag predicts it to some extent. A lowess regression with a restricted span of the smoother shows a peak near zero adaptational lag (Fig. 5D). When mutants appear with effects in the juvenile or adult stage that are slightly advantageous, they will have larger probabilities to get established when the adaptational lag for germination probability in the population is near zero. This adaptational lag can thus be seen as a measure of general adaptability. In Fig. 4F we observed that the lag was smallest in the cooler years near the end of the simulation. Upon inspection, it turns out that the lag is smallest when the total number of recruits per year is largest. This requires, however, phenotypic plasticity of the germination probability. Plastic strategies can exploit the good years to outcompete other strategies more easily by germinating with probability one and don’t need to implement a compromise between good and bad years because of lack of information.

**Figure five.**
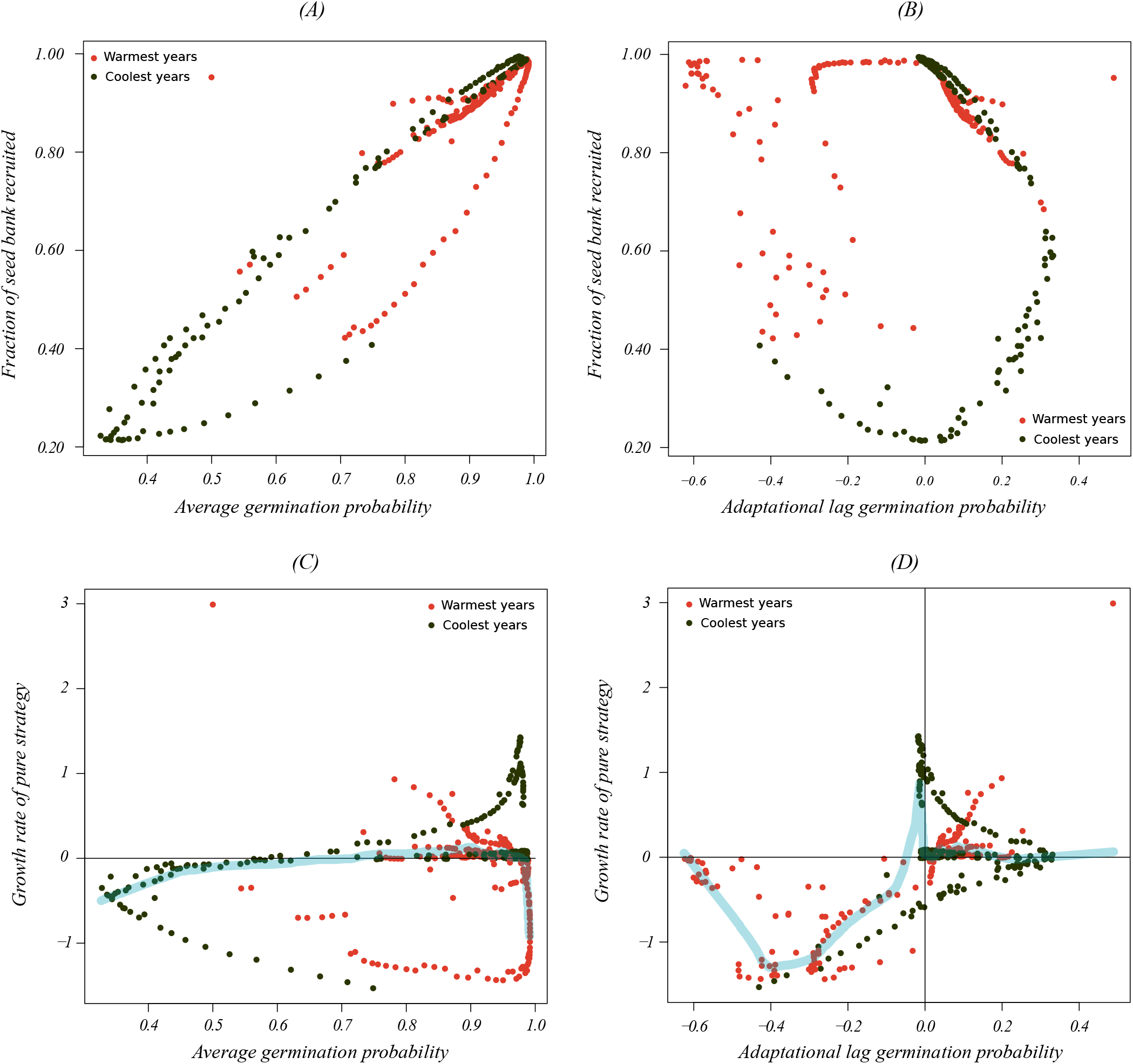
Relationships between demographic statistics and adaptational lag. For the germination probability model with plasticity and maternal effects. (A) and (B) I calculated per year the fraction of the seed bank that was recruited within that year. This statistic is indicative of how fast a population might respond as a whole to selection during the juvenile and adult stages. Panels (A) and (B) show the dependence on average germination probability and adaptational lag (as in Fig. 3F), respectively. (C) and (D) The log per capita number of seeds contributed by a juvenile plant (the growth rate of the pure strategy which always germinates), is a component of the establishment probability of variants. Panel (C) shows its dependence on average germination probability and (D) on the adaptational lag. Lowess regression fits (Cleveland 1981) with a smoother span of 0.1 are added as wide grey lines in panels (C) and (D).

Evolutionary Adaptation and Adaptive Control

There have been repeated calls for an increased use of control theory in ecology, where the most successful applications thus far were in agriculture (Loehle 2006). The objective of control theory is to develop control actions (a “controller”) for dynamical systems such that their outputs follow a desired pattern. Several notions of adaptive control exist (Tyukin 2011), while there is no mention of adaptive control in the review by Loehle (2006) on control theory in ecology. According Walters and Hilborn (1978), adaptive control refers to situations where the best control action of a dynamical system must be established through sequential reassessment of system states and dynamic relationships. This generally requires tuning the controller (Tyukin 2011) in dependence on the state of the environment and the responses of the system. In addition, adaptive control often uses a model-based reference response (MRAC, model reference adaptive control, Frank 2018), and this model can also be updated as part of the tuning or when new knowledge becomes available. MRAC can involve actions with the intention of improving knowledge on the system. Such actions can trade-off or conflict with adaptively managing other targets (Walters & Hilborn 1978) and make adaptive management or control more prone to implementation failure (Allen & Gunderson 2011). The implementation of adaptive control is generally challenging (Anderson & Dehghani 2008). It needs to be determined how concepts of evolutionary adaptation and adaptational lags during climate change relate to adaptive control as defined in control theory and in the context of adaptive management of population dynamical systems.

Next to its relevance for the adaptive management of populations, control theory has been used to derive predictions from modelling for phenotypic traits which can vary continually over time (Frank 2019) and are thus examples of phenotypic plasticity or flexible phenotypes. This line of research can be used to illustrate the principles of adaptive control and to situate it relative to evolutionary adaptation, by gradually incrementing the complexity of a series of trait determination models, where the apparent phenotypic traits are examples of system outputs. A simple partitioning of the contributions of genetic and an environmental input is a common representation of the determination of phenotypes (Fig. 6A&B). The environmental input generates predictable phenotypic consequences in the presence of phenotypic plasticity. There is no control of phenotypes here towards a specific target because all inputs are assumed to have additive and linear effects. Models of phenotypic switches do invoke a sort of controller (Fig. 6B, Leimar et al. 2006), which is usually a non-linear function of an underlying liability. Models of phenotypic plasticity which explicitly assume a controller of plasticity or flexibility exist (Fig.6C, Frank 2019), and classical dynamical models of energy allocation can be interpreted in this manner too (Perrin & Sibly 1993). These controls do not have the recurrent tuning by environment and system states characterizing adaptive control. They can assume that the controller is the result of evolutionary adaptation (Frank 2019) but the control in itself can be non-adaptive in the sense of control theory (compare Fig. 6C with Fig. 6D). On the other hand, experimental studies on the control of gaits or bipedal movement envisage elements of adaptive control in locomotor control: Lam et al. (2006) for example explicitly mention that the nervous system uses an internal model for the control of locomotion.

**Figure six.**
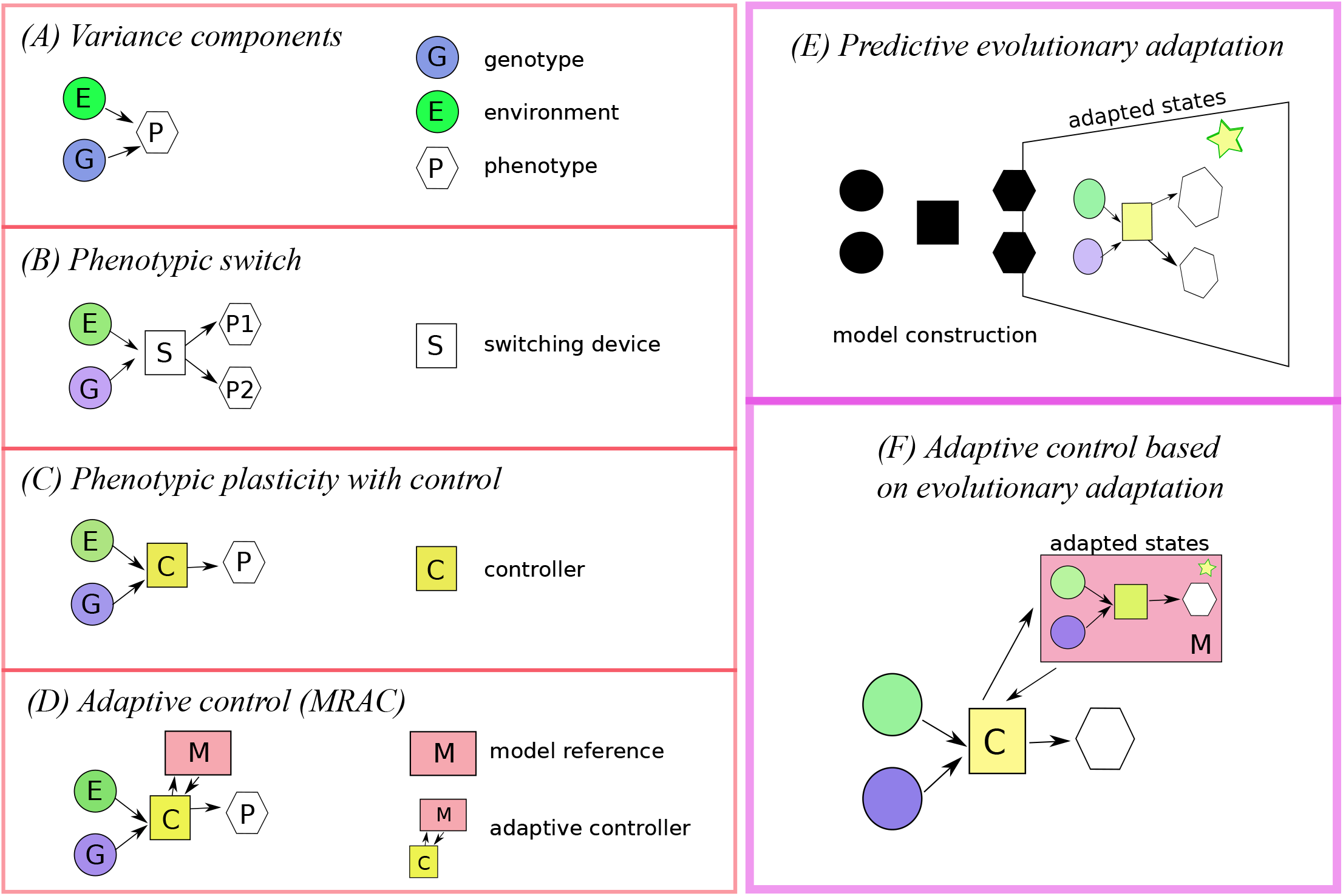
Adaptive control and adapted states. Left. Elaborating modes of trait determination demonstrates that adaptive control shares characteristics with a particular mode of trait determination. Right. Evolutionary adaptation and its integration within an adaptive controller. (A) Phenotypic plasticity (B) Phenotypic switch (C) Plasticity with controller (D) Adaptive control (E) Adaptive dynamics predicts adapted trait values for a trait determination process using a demographic model with environmental feedbacks and mutational variation. (F) These predictions can be used as a model reference in an adaptive control system.

Adaptive control could use a model of evolutionary adaptation (Fig. 6E) to generate a model reference and target of control (Fig. 6F). This would integrate evolutionary adaptation and adaptive control, with the adaptational lag providing the distance from the control target or becoming the quantity to control. This raises the question what an adaptive controller will look like in this case and how to assure that it operates safely without divergence from the target or a general crash of an ecological system. Much remains to be developed. However, there are examples where evolutionary modelling of managed ecosystems led to proposals to control adaptational lags.

#### Adaptive pest and crop management

Studies which considered coevolutionary principles in the adaptive management programs of plant diseases in agriculture, tend to focus on minimizing evolution (Zhan et al. 2015). Pathogens at adapted states are generally undesirable in agricultural practises when properties of the agro-ecology lead to the evolutionary minimization of plant resources available to competing pathogens (Lion & Metz 2018), hence maximization of yield losses. Adaptational lags should then be kept maximal or control should be towards increasing diversifying and disruptive selection weakening selection and imposing trade-offs (Zhan et al. 2015).

Précigout et al. (2020), for example, studied adaptation of the latent period of a plant pathogen to plant traits which depend on fertilization levels. The analysis demonstrated that adaptive pathogen states minimize the amount of healthy leaf canopy, and that it is therefore desirable to maximize the adaptational lag. Preventing adaptation can occur by reducing selection gradients through generally limiting pathogen growth rates (Carolan et al. 2017). Bargués-Ribera and Gokhale (2020) present an example optimizing cumulative yield in dependence on rotation schemes between cash and cover crops. We also start considering the maintenance of pathogen diversity as a useful strategy, if this limits the evolution of pathogen virulence and reduces average pathogen reproduction per host by increasing host diversity (Dutta et al. 2021). However, if adaptation favours a polymorphism of coexisting strains each specializing on a different host plant, avoiding such adaptive genetic polymorphisms can be crucial in heterogeneous landscapes (Précigout 2018). Alternatively, also in plant crops cases are imaginable where adaptational lags need to be maximized, when plant stages are harvested which occur at their lowest densities in adapted states, for example due to asymmetric competition.

It seems easier to imagine examples where adaptational lags in crops should be minimized. Loeuille et al. (2013) use adaptive dynamics modelling to predict consequences of different agricultural practises and landscape designs, presenting an example where adaptation by plants minimizes nutrient loss, therefore maximizes sustainability of the ecosystem.

Adaptation of plants is desirable from this point of view, and adaptive management should aim to minimize the adaptational lag in plants even if this does not necessarily occur by natural selection. van den Bosch et al. (2007) were to my knowledge the first to demonstrate that predictions of adapted states have relevance in agricultural systems and made a point of general relevance: the control strategy should be chosen depending on whether adapted states imply increased or reduced crop damage. van den Bosch et al. (2007) modelled the adaptation of virus strains in response to different disease control strategies and assumed that control would lead to zero adaptational lag. Sanitation by roguing came out as a control strategy where virus evolution leads to more favourable adaptive states, because these reduce virus titre in comparison to virus adaptation to other control strategies.

Among prospects for future research in modelling plant virus epidemiology, Jeger et al. (2018) pointed to the importance of environmental change, of considering evolution but did not propose to integrate both. In the studies above, the target of control was identified, but it remained unclear whether that control in itself would need to occur in an adaptive manner. These studies all assumed that adaptation occurs in a period without environmental change. I conjecture that when environments change gradually and unpredictably to some extent, most control will need to become adaptive.

## Discussion

The term “adaptation” links a cluster of concepts which each are at the core of a subject in climate change research. In this paper, *adaptational lag* is introduced in the context of evolutionary adaptation and its integration into adaptive control theory is proposed. Other related concepts, such as adaptability and adaptive capacity are only minimally discussed here (Nelson et al. 2007, Bateson 2017, Siders 2018, Angeler et al. 2019).

Moving optimum models (Kopp & Matuszewski 2015) generally assume that the adaptive states are optimal phenotypes which follow a prescribed and linear pattern of change. With more involved life histories and selection on different life history components it is known that generally and even without environmental change, fitness is largest for a compromise phenotype which is in many cases not equal to any of the phenotypes maximizing individual life history components (Cotto et al. 2019). Adaptation in complex life histories should be seen in the sense of Levins (1968) where strategies weigh different options they encounter and achieve the best compromise in the face of their constraints. For example, the compromise phenotype of Cotto el al. (2019, their Eqn. 9) then is the adaptive state. When the genetic variance-covariance at the adaptive state would be restricted to what is maintained by selection in the long run, when not assuming linear environmental change in the adapted state or that the adaptational lag equilibrates at a fixed distance from the compromise phenotype, their method would approach the proposal here. Johansson and Jonzén (2012) were the first to consider frequency-dependence within species in their determination of adapted states for bird arrival date. They determined adaptive states but did not consider adaptational lags. Gienapp et al. (2013) inserted adapted states determined for a different model of bird breeding dates as the optimal phenotypes in a standard quantitative genetic model from which adaptational lags were determined. Their study can thus be interpreted as the first where the current approach was implemented.

In principle, the approach to determine adaptational lags based on Adaptive Dynamics can be carried out for any ecological system where the dependencies of demography on environmental state (including feedbacks of population state) are specified and where phenotypic trait determination is known or assumed to work in a particular way such that an evolutionary model can be constructed. In the example of the evolution of germination probability in an annual plant, the adapted states were germination probabilities which depended on the scenario of environmental change. Adaptive Dynamics studies have distinguished various types of adapted states, of which Figure seven gives a non-exhaustive overview. Next to adapted states with single genotypes, with plasticity or not, outcomes such as adaptive genetic polymorphisms can occur (Geritz et al. 1998). For these adapted states, it remains to be worked out how to calculate adaptational lags best, but it has already been proposed that in some cases, maladaptation should be the target here (Précigout 2018).

Johansson (2008) determined lags for one-, two- and three species systems exploiting a resource distribution. These were calculated with respect to the evolutionarily repelling trait matching the resource with the largest carrying capacity, which is opposite the definition of adaptational lag used here based on attracting adapted states. Evolutionary suicide is another possible outcome of adaptation (Gyllenberg & Parvinen 2001), where the adapted state is extinction. Next to the existence of these qualitatively different outcomes, multiple adapted states can co-occur for one environmental setting such that adaptational lags become multivalued. Further outcomes which are not represented in Fig. 7 are coevolutionary adapted states of several species (Débarre et al. 2014), relevant in the agro-ecological context of plants and pests, and adapted states which are cyclic (Dercole & Rinaldi 2008), with a non-equilibrium attractor of the adaptive dynamics.

**Figure seven.**
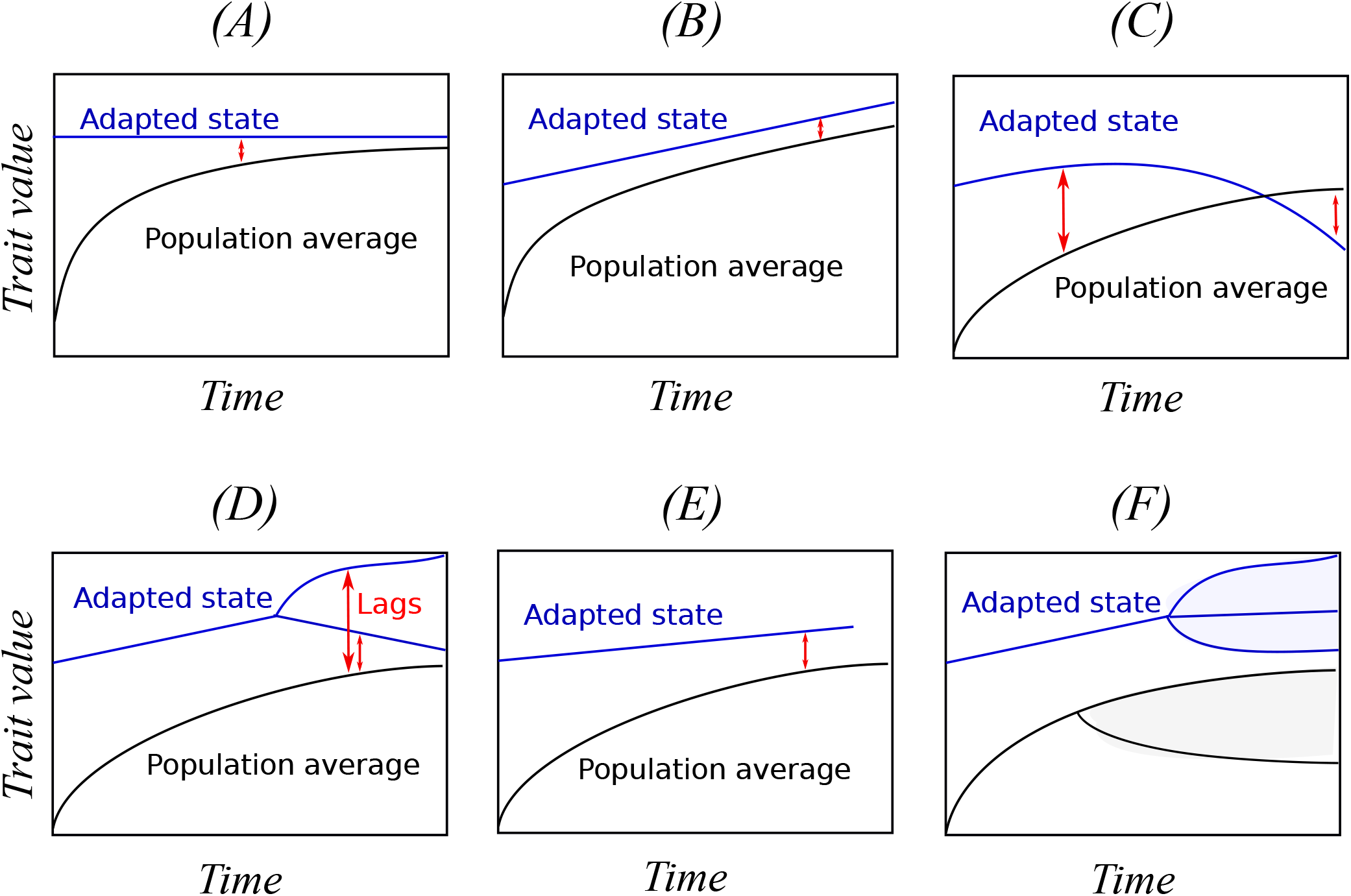
A provisory overview of possible adaptational lag patterns driven by eco-evolutionary dynamics. The *y*-axis represents a scalar phenotypic trait, the *x*-axis time. (A) The population approaches an adapted state. (B) There is convergence to an equilibrium adaptational lag. The population then lags with respect to the adapted state at a fixed distance. (C) The adaptational lag changes sign. The population average trait shows inertia in adaptation, the adaptational lag diverges again. (D) Multiple adapted states can co-occur for the environment specific to one year. The lag then becomes a multi-valued function of time. (E) Evolutionary suicide. The adapted state is extinction. (F) For a range of environmental states, the adapted state is an adaptive polymorphism. The population can then also become phenotypically polymorphic or not.

It was found that adaptational lags can’t be removed for boundary adaptive strategies if any non-adaptive genotypic variance remains present, and in general, extreme strategies face a variance load (Lande & Shannon 1996) with any individual phenotypic variance present. If we extend this idea to human socio-economic evolution, climate adaptation strategies which are extreme and therefore boundary strategies, will never be reached as long as variation in our population remains. Beckage et al. (2018) modelled the feedback between human behaviour and projected climate change. Nudging the perceived risk of climate change was proposed as a mitigation policy response, but whether behavioural responses will be adaptive was not considered. In evolutionary ecology, cue perception and associated response plasticity are often studied as adaptations and even if we refuse to compare our own adaptation to an evolutionary response, it might be useful to try to determine the adapted states of what we aim to implement. Despite the differences between evolutionary adaptation and adaptive control, they can be modelled jointly. In the adaptation policy context, clear metrics of adaptation in comparison to mitigation seem to lack (Michaelowa & Stadelmann 2018). Using an adaptive control perspective in adaptation policies naturally leads to relevant metrics: they are the distances to the model reference which the controller assesses and uses for its updates. The examples of control and evolutionary adaptation in agro-ecologies can be used to model and determine strategies for adaptive management during environmental change and can lead to strategies for developing appropriate controllers.

From the simulation results, it turned out that adaptational lags might be partial predictors of the invasion success of mutants in other life history traits than germination. If we develop our understanding of adaptational lags further, it could be of use in conservation biology and agro-ecology as partial measures of adaptability. For extant populations, would it be possible to estimate adaptational lag and adaptive state approximately with limited modelling? In models with pre-defined adaptive states, the difference between the population average of a trait and its adaptive state is a commonly used measure of the lag (Kopp & Matuszewski 2015) and in the vicinity of the adapted state, it can be determined from slope and curvature of fitness surfaces (Estes & Arnold 2007). In the current environment, selection optima can be estimated using data on the dependency of the number of offspring on phenotype, and the changes in optima over successive years can be tracked or projected. As a first approximation, we could use these as proxies. However, additional modelling and data analysis should be carried out alongside such estimation to determine whether the approximation is warranted and remains useful during environmental change. In other words, we should implement an adaptive control strategy.

## Acknowledgements

This manuscript has taken years to develop due to my own environmental changes, and has been adapted a few times. It has benefited from discussions with Marcel Visser, Luis Chevin and Marc Girondot and from numerous invitations for seminars where I could gauge my ideas on an audience.

## Methods

Adaptive Dynamics approximations describe evolutionary processes where it is assumed that characters exist which are faithfully replicated in offspring (Metz et al. 1996), except for rare mutational changes. Additionally, mutations are often assumed to have small effects at the individual phenotypic level (Dieckmann and Law 1996, Champagnat et al. 2002).

Characteristic is that evolving populations can feed back onto selective pressures through the selective environments they create. However, it is assumed that external environmental fluctuations that act as input are stationary (Ripa and Dieckmann 2003, Geritz et al. 1998) and that the rarity of mutational changes makes the so-called resident environments stationary too. Based on all these assumptions, a deterministic dynamic system can be defined, referred to as the canonical equation of AD (Dieckmann and Law 1996). Not assuming small mutations implies that evolutionary trait substitution sequences are studied (Champagnat et al. 2002) where in each substitution a resident genotype is replaced by a mutant with positive invasion fitness. Convergence and stability of singular points is then based on geometric arguments, not directly deriving from a deterministic dynamical system. Evolutionarily singular points (candidate adapted states) can be CSS endpoints of trait substitution sequences or the canonical equation of Adaptive Dynamics when conditions on second order partial derivatives of invasion fitness confirm convergence stability and non-invasibility (Geritz et al. 1998). CSS’s and their properties are relatively independent of genetic makeup when adapted states contain one phenotype (Van Dooren 2006). Systematic phenotypic advantages for heterozygotes are not expected for most smooth genotype-phenotype maps (Van Dooren 2000).

Simulation code for R used to study the germination probability model is available on request. I simulated populations of selfing plants according the model described in the main text. For the simulation with genotypically determined germination probabilities, trait values were discretized from zero to one in steps of 0.02. The initial state of the population was a uniform distribution of individuals over all trait values, with densities summing up to value one half. The mutation probability was set at 0.1 per individual and mutants occupied trait bins adjacent to their parental trait value. To determine adapted states for this model version with genotypic germination probabilities only, evolutionary trait substitution sequences were simulated with trait step sizes of 0.1 starting from germination probability one and trait boundaries at values 0.1 and one. For the forward simulations with plasticity and maternal effects, trait values were discretized at integer values from -5 to +5 for each trait *g, b, m* resulting in 1331 trait combinations which occurred. The mutation rate was set at 0.1 per individual. Mutant individuals were uniformly distributed over all bins adjacent to their parental trait combination and within the range of trait values allowed. The initial state of the population was in each forward simulation a uniform distribution over all trait combinations, with densities summing up to value one. For the evolutionary invasion analysis used to determine adapted states, evolutionary trait substitution sequences were simulated with trait step sizes of 0.1 and starting from zero trait values for *g, b* and *m*. At each substitution, simultaneous trait changes were made in all traits with a direction of positive invasion fitness. Trait boundaries were fixed at -5, 5 for each trait.

## Notes

### Competing Interest Statement

The authors have declared no competing interest.

### Summary of Updates

The section on adaptive control has been reworked.

## References

Abrams, PA (2005) Adaptive Dynamics’ vs.’adaptive dynamics. J. evol. Biol. 18: 1162–1165.

Allen, CR, Gunderson, LH (2011) Pathology and failure in the design and implementation of adaptive management. J Environ Manage. 92: 1379–84.

Amundson, R (1996) Historical development of the concept of adaptation. In: Adaptation, eds. Rose, MR, Lauder, GV. Academic Press, New York, pp. 11–54.

Anderson, B, Dehghani, A (2008) Challenges of adaptive control-past, permanent and future. Annu. Rev. Control. 32: 123–135.

Angeler, DG, Fried-Petersen, H, Allen, CR, Garmestani, A, Twidwell, D, Birgé, HE et al. (2019) Adaptive capacity in ecosystems. Adv Ecol Res. 60: 1–24.

Bargués-Ribera, M, Gokhale, CS (2020) Eco-evolutionary agriculture: Host-pathogen dynamics in crop rotations. PLoS Comput. Biol. 16: e1007546

Bateson, P (2017) Adaptability and evolution. Interface Focus. 7: 20160126.

Beckage, B, Gross, LJ, Lacasse, K, Carr, E, Metcalf, SS, Winter, JM et al. (2018) Linking models of human behaviour and climate alters projected climate change. Nat. Clim. Change 8: 79.

Botero, CA, Weissing, FJ, Wright, J, Rubenstein DR (2015) Evolutionary tipping points in the capacity to adapt to environmental change. Proc. Natl. Acad. Sci. U.S.A. 112: 184–189.

Bulmer, MG (1984) Delayed germination of seeds: Cohen’s model revisited. Theor. Pop. Biol. 26: 367–377.

Carolan, K, Helps, J, van den Berg, F. Bain, R, Paveley, N, van den Bosch, F (2017) Extending the durability of cultivar resistance by limiting epidemic growth rates. Proc. Royal Soc. B 284: 20170828.

Champagnat, N, Ferrière, R, Ben Arous, R (2001) The canonical equation of adaptive dynamics: a mathematical view. Selection 2: 73–84.

Cleveland, WS (1981) LOWESS: A program for smoothing scatterplots by robust locally weighted regression. Am. Stat. 35, 54.

Cohen, D (1966) Optimizing reproduction in a randomly varying environment. J. Theor. Biol. 12: 110–129.

Collins, MR, Knutti, J, Arblaster, J-L, Dufresne, T, Fichefet, P, Friedlingstein, X, et al. (2013) Long-term climate change: Projections, commitments and irreversibility. In: Climate Change 2013: The Physical Science Basis. Contribution of Working Group I to the Fifth Assessment Report of the Intergovernmental Panel on Climate Change. Eds. Stocker, TF, Qin, D, Plattner, G-K, Tignor, M, Allen, SK, Doschung, J, Nauels, A, Xia, Y, Bex, V, Midgley PM. Cambridge University Press, pp. 1029–1136.

Cotto, O, Sandell, L, Chevin, LM, Ronce, O (2019) Maladaptive shifts in life history in a changing environment. Am. Nat. 194: 558–573.

Coulson, T (2020) Equilibrium, the new dirty word of ecology. Authorea. Available at https://doi.org/10.1111/ele.13629.

Débarre, F, Nuismer, SL, Doebeli, M (2014) Multidimensional (co) evolutionary stability. Am. Nat. 184: 158–171.

Dercole, F, Rinaldi, S (2008) Analysis of Evolutionary Processes: The Adaptive Dynamics Approach and its Applications. Princeton University Press, Princeton New Jersey.

Dieckmann, U (1997) Can adaptive dynamics invade? Trends Ecol Evol. 12:128–31.

Dieckmann, U, Law, R (1996) The dynamical theory of coevolution: a derivation from stochastic ecological processes. J Math Biol. 34: 579–612.

Dutta, A, Croll, D, McDonald, BA, Barrett, LG (2021) Maintenance of variation in virulence and reproduction in populations of an agricultural plant pathogen. Evol. Appl. 14: 335–47.

Ellner, S (1997) You bet your life: life-history strategies in fluctuating environments. In: Case Studies in Mathematical Modeling: Ecology, Physiology, and Cell Biology. Eds. Othmer, HG, Adler, FR, Lewis, MA, Dallon, JC. Prentice-Hall, New Jersey, pp. 3–24.

Eshel, I (1983) Evolutionary and continuous stability, J. theor. Biol. 103: 99–111.

Estes, S, Arnold, SJ (2007) Resolving the paradox of stasis: models with stabilizing selection explain evolutionary divergence on all timescales. Am. Nat. 169: 227–44.

Ferrière, R, Gatto, M (1995) Lyapunov exponents and the mathematics of invasion in oscillatory or chaotic populations. Theor. Pop. Biol. 48: 126–171.

Ferrière R, Legendre S (2013) Eco-evolutionary feedbacks, adaptive dynamics and evolutionary rescue theory. Phil. Trans R Soc B 368:20120081.

Fisher, RA (1930). The Genetical Theory of Natural Selection. Clarendon Press, Oxford.

Frank, SA (2018) Control Theory Tutorial: Basic Concepts Illustrated by Software Examples. Springer, Cham, Switzerland.

Frank SA (2019) Evolutionary design of regulatory control. I. A robust control theory analysis of tradeoffs. J. Theor. Biol. 463: 121–137

Geritz, SAH, Meszéna, G, Metz, JAJ (1998) Evolutionarily singular strategies and the adaptive growth and branching of the evolutionary tree. Evol. Ecol. 12:.35–57.

Gienapp, P, Merilä, J (2018) Evolutionary responses to climate change. In: Encyclopedia of the Anthropocene, eds. DellaSalla, D, Goldstein, M. Elsevier, Oxford, pp. 51–59.

Gienapp, P, Lof, M, Reed, TE, McNamara, J, Verhulst, S, Visser, ME (2013) Predicting demographically sustainable rates of adaptation: Can great tit breeding time keep pace with climate change? Philos. Trans. R. Soc. Lond. B Biol. Sci. 368: 20120289.

Gyllenberg, M, Parvinen, K (2001) Necessary and sufficient conditions for evolutionary suicide. Bull. Math. Biol. 63: 981–993.

Haldane, JBS (1927) A mathematical theory of natural and artificial selection, Part V: selection and mutation. Math. Proc. Camb. Philos. Soc. 23: 838–844.

Johansson, J (2008) Evolutionary responses to environmental changes: how does competition affect adaptation? Evolution 62: 421–435.

Johansson, J, Jonzén, N (2012) Game theory sheds new light on ecological responses to current climate change when phenology is historically mismatched. Ecol. Lett. 15: 881–888.

Jeger, MJ, Madden, LV, van den Bosch, F (2018) Plant virus epidemiology: applications and prospects for mathematical modeling and analysis to improve understanding and disease control. Plant Dis. 102: 837–854.

Kane, S, Shogren, JF (2000) Linking adaptation and mitigation in climate change policy. In: Societal Adaptation to Climate Variability and Change. Eds. Kane, SM, Yohe, GW. Springer, Dordrecht, pp. 75–102.

Kopp, M, Matuszewski, S (2014) Rapid evolution of quantitative traits: theoretical perspectives. Evol. Appl. 7: 169–91.

Kuznetsov, YA (2004) Elements of Applied Bifurcation Theory (Vol. 112), Third Edition. Springer Science & Business Media, New York.

Lam T, Anderschitz M, Dietz V. (2006) Contribution of feedback and feedforward strategies to locomotor adaptations. J Neurophysiol. 95: 766–73.

Lande, R, Shannon, S (1996) The role of genetic variation in adaptation and population persistence in a changing environment. Evolution 50: 434–437.

Leimar, O (2009) Multidimensional convergence stability. Evol. Ecol. Res. 11: 191–208.

Leimar, O, Hammerstein, P, Van Dooren, TJM (2006) A new perspective on developmental plasticity and the principles of adaptive morph determination. Am Nat. 167: 367–76.

Levins, R (1968) Evolution in Changing Environments. Some Theoretical Explorations. Princeton University Press, Princeton New Jersey.

Lion S, Metz JAJ. (2018) Beyond R0 maximisation: on pathogen evolution and environmental dimensions. Trends Ecol Evol. 33: 458–473.

Loehle, C (2006) Control theory and the management of ecosystems. J. Appl. Ecol. 957–966.

Loeuille, N, Barot, S, Georgelin, E, Kylafis, G, Lavigne, C (2013). Eco-evolutionary dynamics of agricultural networks: implications for sustainable management. Adv. Ecol. Res 49: 339–435.

Lynch, M, Gabriel, W, Wood, AM (1991) Adaptive and demographic responses of plankton populations to environmental change. Limnol. Oceanogr. 36: 1301–1312.

Merilä, J, Hendry, AP (2014) Climate change, adaptation, and phenotypic plasticity: the problem and the evidence. Evol. Appl. 7: 1–14.

Metz, JAJ, Nisbet, RM, Geritz, SAH (1992) How should we define ‘fitness’ for general ecological scenarios? Trends Ecol. Evol. 7: 198–202.

Metz, JAJ, Geritz, SAH, Meszena, G, Jacobs, FJA, van Heerwaarden, JS (1996) Adaptive dynamics: A geometrical study of the consequences of nearly faithful reproduction. In: Stochastic and Spatial Structures of Dynamical Systems. Eds. Van Strien, SJ, Verduyn Lunel, SM. North-Holland, Amsterdam, pp. 183–231.

Michaelowa, A, Stadelmann, M (2018) Development of universal metrics for adaptation effectiveness. In: Adaptation metrics: perspectives on measuring, aggregating and comparing adaptation results. Eds. Christiansen, L, Martinez, G Naswa, p. UNEP DTU Partnership, Copenhagen, pp. 63–72.

Nelson, DR, Adger, WN, Brown, K (2007) Adaptation to environmental change: contributions of a resilience framework. Annu. Rev. Environ. Resour. 21: 32:395–419.

Newman, JA, Anand, M, Henry, HAL, Hunt, S, Gedalof, Z (2011) Climate change biology. CABI, Cambridge, MA.

Ooi, M (2012). Seed bank persistence and climate change. Seed Science Res. 22: S53–S60.

Otto, SP, Day, T (2007) Evolutionary invasion analysis. In: A Biologist’s Guide to Mathematical Modeling in Ecology and Evolution. Princeton University Press, New Jersey, pp. 454–502.

Peischl, S, Kirkpatrick, M (2012) Establishment of new mutations in changing environments. Genetics 191: 895–906.

Perrin, N, Sibly, RM (1993) Dynamic models of energy allocation and investment. Annu. Rev. Ecol. Evol. Syst. 24: 379–410.

Précigout, PA. (2018) Epidemiology and evolution of fungal foliar pathogens in the face of changes in crop fertilization : application of evolutionary-ecological theory to crop epidemiology. PhD thesis Université Sorbonne Paris Cité. Available at: https://tel.archives-ouvertes.fr/tel-02333571

Précigout PA, Robert C, Claessen D (2020) Adaptation of biotrophic leaf pathogens to fertilization-mediated changes in plant traits: a comparison of the optimization principle to invasion fitness. Phytopathology 110: 1039–1048.

Rand, DA, Wilson, HB, McGlade, JM (1994) Dynamics and evolution: evolutionarily stable attractors, invasion exponents and phenotype dynamics. Phil. Trans. R. Soc. B 343: 261–283.

Ripa, J, Dieckmann, U (2013) Mutant invasions and adaptive dynamics in variable environments. Evolution 67: 1279–1290.

Sabelis, MW, Metz, JAJ (2002) Taking stock: relating theory to experiment. In: Adaptive Dynamics of Infectious Diseases: in Pursuit of Virulence Management. Eds. Dieckmann, U, Metz, JAJ, Sabelis, MW, Sigmund, K. Cambridge Series in Adaptive Dynamics, Cambridge University Press, Cambridge UK.

Schipper, ELF (2006) Conceptual history of adaptation in the UNFCCC process. Rev. Eur. Comp. Int. Environ. Law 15: 82–92.

Siders, AR (2019) Adaptive capacity to climate change: A synthesis of concepts, methods, and findings in a fragmented field. Wiley Interdiscip. Rev. Clim. Change 10: e573.

Stearns, SC (2000) Life history evolution: successes, limitations, and prospects. Naturwissenschaften 87: 476–486.

Teplitsky, C, Tarka, M, Møller, AP, Nakagawa, S, Balbontín, J, Burke, TA, et al. (2014) Assessing multivariate constraints to evolution across ten long-term avian studies. Plos One E 9: e90444.

Tuljapurkar, S (1986) Demography in stochastic environments. II. Growth and convergence rates. J Math Biol. 24: 569–81.

Tyukin, I (2011) Adaptation in dynamical systems. Cambridge University Press, New York.

van den Bosch, F, Jeger, MJ, Gilligan, CA (2007) Disease control and its selection for damaging plant virus strains in vegetatively propagated staple food crops; a theoretical assessment. Proc Biol Sci. 274: 11–18.

Van Dooren, TJM (2000) The evolutionary dynamics of direct phenotypic overdominance: emergence possible, loss probable. Evolution 54: 1899–1914.

Van Dooren, TJM (2006) Protected polymorphism and evolutionary stability in pleiotropic models with trait-specific dominance. Evolution 60: 1991–2003.

Walsh, B, Lynch, M (2018) Evolution and selection of quantitative traits. Oxford University Press, Oxford.

Walters, CJ, Hilborn, R (1978) Ecological optimization and adaptive management. Annu. Rev. Ecol. Evol. Syst. 9: 157–188

Zhan J, Thrall PH, Papaïx J, Xie L, Burdon JJ (2015) Playing on a pathogen’s weakness: using evolution to guide sustainable plant disease control strategies. Annu Rev Phytopathol. 53: 19–43.

